# Genomic analysis of extended-spectrum β-lactamase-producing *Enterobacter kobei* ST691 strain harbouring *mcr*-10.1 isolated in Yaoundé, Cameroon

**DOI:** 10.64898/2025.12.28.696731

**Authors:** Raspail Carrel Founou, Luria Leslie Founou, Erkison Ewomazino Odih, Anderson O. Oaikhena, Iruka N. Okeke

## Abstract

**Objective:** *Enterobacter species*. are opportunistic pathogens commonly responsible for serious, difficult-to-treat hospital-acquired infections. Extended-spectrum β-lactamase (ESBL)-producing and colistin-resistant *Enterobacterales* are increasingly implicated in human and animal infections worldwide. Here we report a first detection of colistin-resistant ESBL-producing *E. kobei* strain belonging to ST691 and harbouring *mcr*-10.1.

**Methods:** This strain was isolated from the faecal sample of a two-year old child aged, who was diagnosed with gastroenteritis in Yaoundé, Cameroon. This ESBL-*E. kobei* ST691 genome was sequenced using Illumina Miseq (Illumina, San Diego, CA, USA). Mobile genetic elements and antibiotic resistance genes were predicted using AMRFinderPlus, ARIBA and the VFDB and PlasmidFinder databases respectively.

**Results:** This strain exhibited phenotypic resistance to numerous antibiotics belonging to penicillin, third generation of cephalosporin and carbapenem families. However, it was susceptible to aminoglycoside and fluoroquinolone. Genome analysis reveals a length of 4 626 300 bp, and N50 of 143 731, GC content of 54.9%. Genes conferring resistance to β-lactams (*bla*_ACT-9_), polymyxin (*mcr-*10.1) and phenicol/quinolone (*oqx*A,B) were detected. Mobile genetic elements including plasmid replicon type [IncFIB(pECLA) and IncFII(pECLA)] and the IS1, IS110, ISEc34, IS1222SC, IS66, IS630, IS3, IS26 insertion sequences were also detected.

**Conclusion:** We report the first colistin-resistant ESBL-producing *E. kobei* isolated from a two-year old child. It is a high priority potential pathogen showing resistance to last-resort antimicrobials to which there is little access in Cameroon. This underscores the necessity to strengthen genomic surveillance, antimicrobial stewardship and infection prevention and control as well as to heighten awareness of the threat posed by resistant bacteria.

## 1. Introduction

*Enterobacter species*. are opportunistic pathogens responsible for difficult-to-treat hospital-acquired infections, including UTIs, skin infections and bacteraemia [1]. They overproduce chromosomally encoded cephalosporinase (cAmpC) associated with the permeability reduction of the outer membrane which may confer reduced susceptibility to carbapenem [2]. The emergence of carbapenem-and colistin-resistant *E. kobei* limits therapeutic options [1]. This strain is increasingly important pathogens implicated in human and animal infections [1,3] and carbapenem-resistant *Enterobacter* spp. are listed by the World Health Organisation (WHO) as critical priority pathogens for which new antimicrobials need to be developed [4].

We report here the first description of a colistin-resistant and extended-spectrum beta-lactamase (ESBL)-producing *Enterobacter kobei* strain (PR13) harbouring *bla*_ACT-9_ and *mcr*-10.1. It was isolated from the faecal sample of a two year old child aged, who was diagnosed with gastroenteritis in Cameroon. Multi-drug resistance of this isolate prompted whole genome sequencing with the goal of understanding the genetic basis of the observed resistance phenotype.

## 2. Materials and methods

During a four-month period from July to October 2020, clinical specimens at two health facilities were screened for ESBL-producing *Enterobacterales*. The specimens screened included wound swabs, urine, uro-vaginal swabs, and stool samples. Specimens were collected and transported to the microbiology laboratory of the Research Institute of the Centre of Expertise and Biological Diagnostic of Cameroon (CEDBCAM-RI) within two hours of collection. Stool samples were cultured on Mac Conkey agar supplemented with crystal violet. ESBL screening was done using CHROMagar™ ESBL according to the manufacturer’s instructions. Identification was performed using biochemical profile with API 20E and confirmed with VITEK 2 system and MALDI-TOF. Antimicrobial susceptibility testing was performed using the Kirby-Bauer disc diffusion method and minimum inhibitory concentrations were obtained with the Vitek® 2 System using Gram Negative Susceptibility card (AST-N255) (BioMérieux, Marcy l’Etoile, France). The European Committee on Antimicrobial Susceptibility testing guidelines 2019 was used for interpretation of the results and *E. coli* ATCC 25922 and *K. pneumoniae* ATCC700603 were used as controls.

Genomic DNA (gDNA) was extracted using the Wizard® Genomic DNA Purification Kit (Promega, Wisconsin, USA) according to the manufacturer’s instructions. NanoDrop spectrophotometry and fluorometric analysis (Qubit®) were used to verify the integrity and purity of the gDNA. Double-stranded DNA libraries were prepared with the NEBNext Ultra II FS DNA Library Prep Kit for Illumina (New England Biolabs, Massachusetts, USA) following Global Health Research Unit (GHRU) in-house protocols. Library concentration and fragment distribution were analysed with Qubit dsDNA High Sensitivity Assay Kit (Thermo Fisher Scientific, Massachusetts, USA) on a Qubitflex fluorometer and Agilent High Sensitivity DNA Kit (Agilent Technologies, California, USA) on a bioanalyser. Libraries were then denatured and sequenced on an Illumina Miseq using paired-end 2 by 150 bp reads (Illumina, San Diego, CA, USA). The sequenced reads were assembled using the GHRU assembly pipeline with default parameters and genome quality cut-offs (https://www.protocols.io/view/ghru-genomic-surveillance-of-antimicrobial-resista-bpn6mmhe). The genome was annotated using the National Center for Biotechnology Information’s Prokaryotic Genome Automated Pipeline and antimicrobial resistance genes were identified using AMRFinderPlus v3.10.24. Plasmids were detected using PlasmidFinder 2.1 (http://cge.cbs.dtu.dk/services/PlasmidFinder/) and the Comprehensive Antibiotic Resistance Database (CARD) respectively.

## 3. Results

Among 49 non-duplicate ESBL-*Enterobacterales* from clinical specimens (endocervical swab, blood, urine, feces and throat swab), three were *Enterobacter* spp. including *Enterobacter kobei* (stool), *Enterobacter hormaechei* (wound swab) and *Enterobacter asburiae* (uro-genital swab). The sole *E. kobei* from a two-year old child with the gastroenteritis symptoms (stomach-ache, vomiting and diarrhoea), showed resistance to amoxicillin/clavulanic acid, ticarcillin/clavulanic acid, cefuroxime cefotaxime, ceftazidime chloramphenicol and meropenem. However, it was susceptible to imipenem, ertapenem, amikacin, gentamicin, ofloxacin, ciprofloxacin, levofloxacin and sparfloxacin.

The genome of *E. kobei* isolate has a genome length of 4 626 300 bp, an N50 of 143 731, GC content of 54.9% and was deposited in the European Nucleotide Archive (https://www.ebi.ac.uk/ena) with accession No. **ERR10862935**. It was assigned to the sequence type (ST) 691. Genes conferring resistance to β-lactams (*bla*_ACT-9_), polymyxin (*mcr-*10.1) and phenicol/quinolone (*oqx*A,B). Two plasmid replicon types, namely IncFIB(pECLA) (Accession no. **CP001919**; 100% identity, contig 51) and IncFII(pECLA) (Accession no. **CP001919**; 99.6% identity, contig 56) were detected in the isolate’s genome, as were IS1, IS110, ISEc34, IS1222SC, IS66, IS630, IS3, IS26, and cn_4541_ISSgsp1 insertion sequences.

VirulenceFinder revealed the *mrkABCDF* fimbrial operon promoting adherence and biofilm formation among *Enterobacterales*. Interestingly, a BLAST analysis of the *mrkABCDF* contig revealed that it is identical to a 2.5kb region of an *E. coli* IncFII plasmid (accession: **CP088133.1**) that also carries *mcr-*10.

The Figure 1 showed selected *E. kobei* genomes in Pathogenwatch (https://pathogen.watch) belonging to distinct sequence types and isolated from distinct countries and constructed a maximum likelihood phylogeny. This isolate was closely related to an *E. kobei*, recovered from a human (USA) in 2003 (https://microreact.org/project/8xhqAsqugKT6RmTAHHNFEW-ekobeiyaounde) [5].

**Figure 1:**
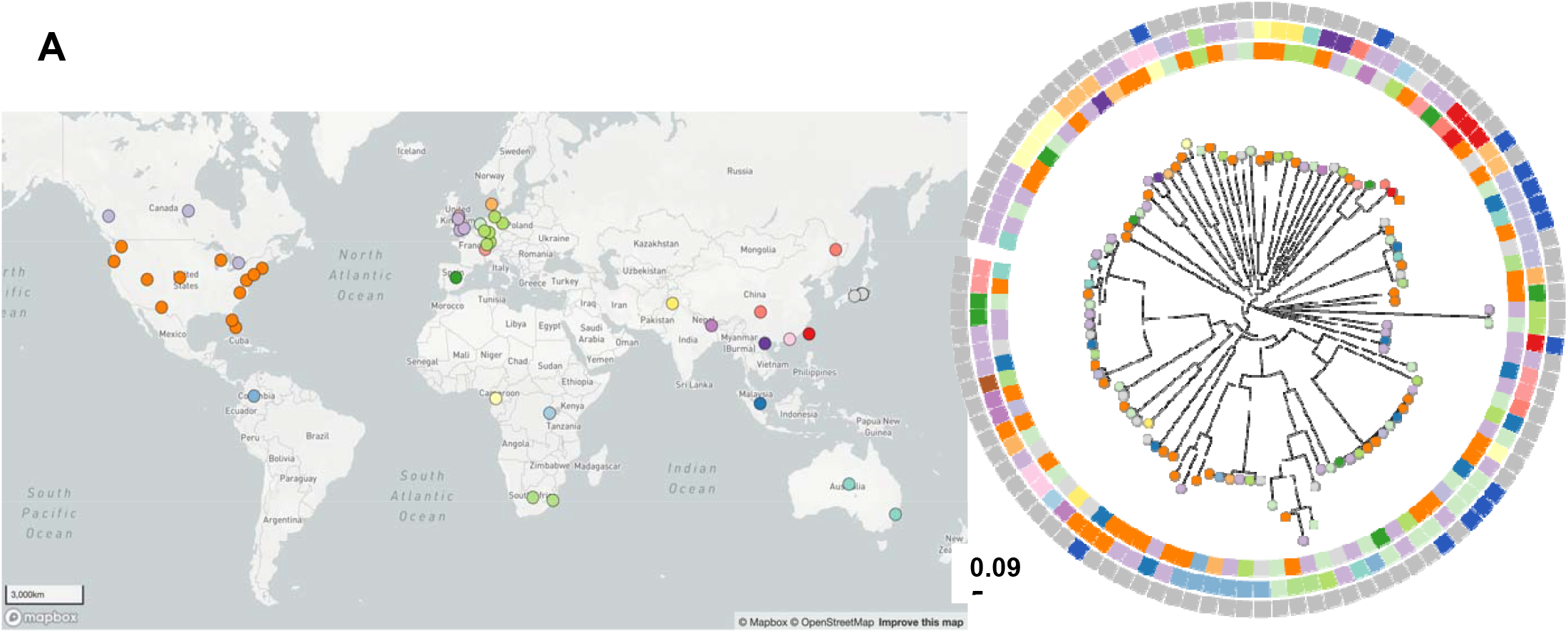
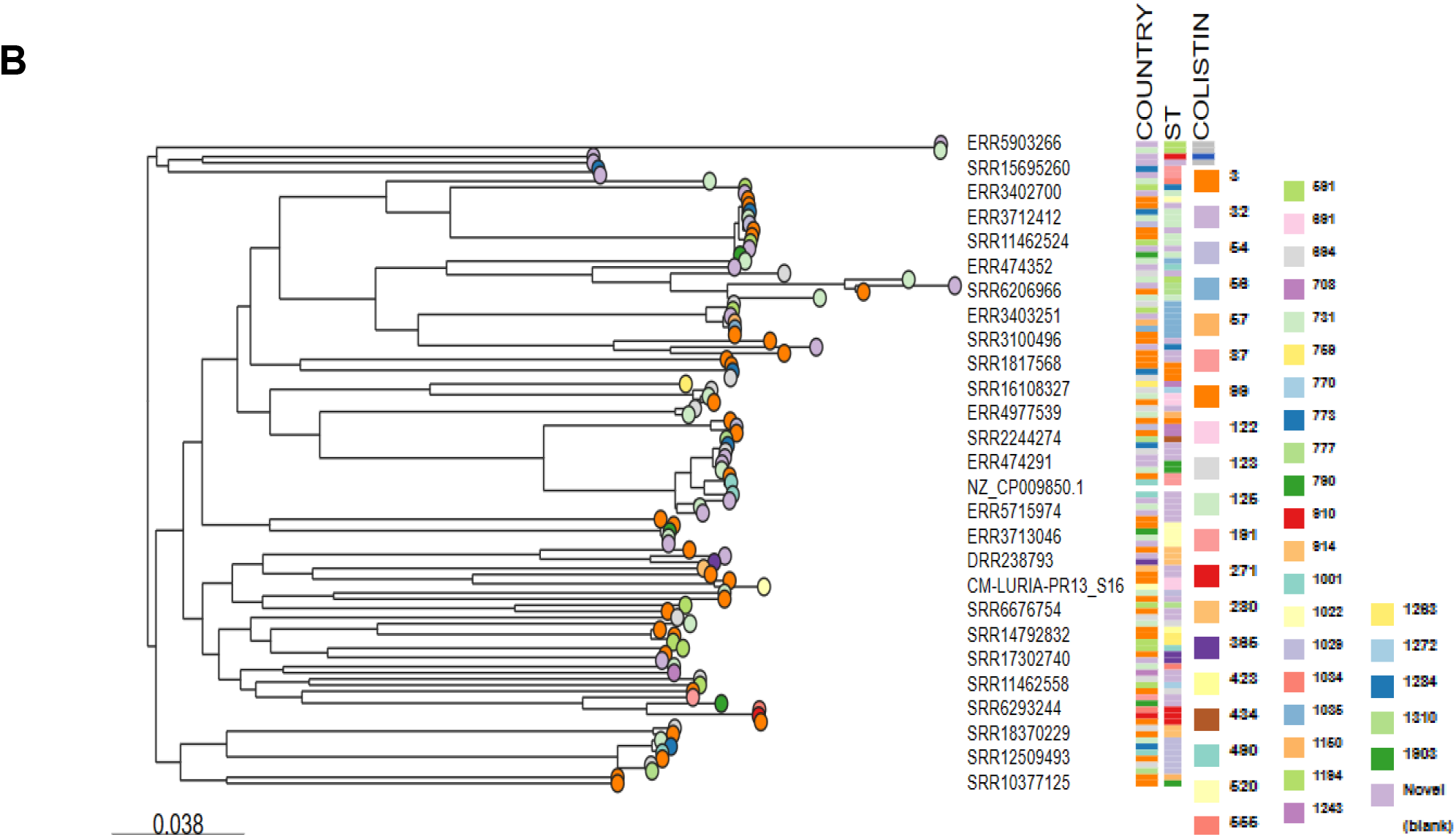
Phylogeny based on core genome multi-locus sequence typing genes of 27 *E. kobei* genomes. The following information is provided for each isolate: name/reference, ST types (STs), country and colistin resistance. STs are highlighted as indicated in the legend and isolate name’s present in the study is CM-LURIA-PR13_S16.

## 4. Discussion

We described here the first report of a colistin-resistant ESBL-producing *E. kobei* stool harbouring *mcr*-10.1 and it was isolated from a child of two-year old with gastro-enteritis that unlikely to have been exposed to colistin. Mobile colistin resistance (*mcr*) genes confer resistant to a last-line antibacterial therapy and are increasingly detected in isolates from humans, animals and the environment worldwide [6]. Recent reports from the Republic of Korea showed that clinical *E. kobei* co-harbouring *mcr*-4.3, *mcr*-9 and *bla*_ACT-64_ were implicated in infections among elderly patients attending emergency department [6]. It has also been detected in a Franciscana dolphin in Brazil, chickens and exposed workers in Asia [6-7]. *E. kobei* bearing *bla*_ACT-9_ and *mcr*-9 have recently been reported from wastewater in South Africa [8]. The strain carrying the gene had mobile element signatures are indicative of transmission potential and the isolation of this strain points to a human faecal reservoir of a priority pathogen.

## 5. Conclusion

Beta-lactam antimicrobials are a mainstay for addressing bacterial pathogens in Cameroon and colistin is considered as a last resort treatment option among patients infected with resistant strains. The detection of *Enterobacter kobei* bearing resistance genes to both classes of antimicrobials emphasizes the need to reinforce antimicrobial stewardship programmes, heighten awareness among physicians and general population, strengthen and implement strict infection prevention and control measures in healthcare settings to curb the dissemination of colistin-resistant ESBL-*E. kobei*.

## Acknowledgements

We are grateful to Faith I. Oni and Odion O. Ikhimiukor for technical assistance, Jola-Ade J Ajiboye for administrative support and the NCBI Genbank submission staff for help with genome upload, decontamination and deposition procedures. We thank other members of the SEQAFRICA consortium, in particular Rene Hendricksen and Pernille Nilsson, for helpful discussions and program oversight.

## Transparency Declarations

The authors declare that they have no known competing financial interests or personal relationships that could have appeared to influence the work reported in this paper.

## Ethical approval

Ethical approval was obtained from the institutional ethics of Research in Human Health **(CEIRSH)** (**No. 2020/020804/CEIRSH/ESS/MIM)**. Permission to conduct the research was also granted from the Head Department of Health. The study was conducted in accordance with the declaration of Helsinki. In addition, the research authorizations of the various healthcare structures have been granted. We had assured the confidentiality of the patient information’s and only the principal investigator had this information. Moreover, this information was anonymized and all isolates were stored for further research.

## Author contributions

**RCF** co-conceptualized the study, undertook sample collection, microbiological laboratory and data analyses, prepared tables and figures, interpreted results, contributed to bioinformatics analysis, and drafted the manuscript. **LLF** undertook sample collection, microbiological laboratory analyses, contributed to bioinformatics analysis and vetted the results. **EEO** performed whole genome sequencing analysis, prepared tables and figures, interpreted results and edited the manuscript. **AOO** set up and oversaw the sequencing workflow and validated the identity and antimicrobial susceptibility of the isolate. **INO** co-conceptualized the study and undertook critical revision of the manuscript. All authors read and approve the final manuscript.

